# Multi-Branch-CNN: classification of ion channel interacting peptides using parallel convolutional neural networks

**DOI:** 10.1101/2021.11.13.468342

**Authors:** Jielu Yan, Bob Zhang, Mingliang Zhou, Hang Fai Kwok, Shirley W. I. Siu

## Abstract

Ligand peptides that have high affinity for ion channels are critical for regulating ion flux across the plasma membrane. These peptides are now being considered as potential drug candidates for many diseases, such as cardiovascular disease and cancers. There are several studies to identify ion channel interacting peptides computationally, but, to the best of our knowledge, none of them published available tools for prediction. To provide a solution, we present Multi-branch-CNN, a parallel convolutional neural networks (CNNs) method for identifying three types of ion channel peptide binders (sodium, potassium, and calcium). Our experiment shows that the Multi-Branch-CNN method performs comparably to thirteen traditional ML algorithms (TML13) on the test sets of three ion channels. To evaluate the predictive power of our method with respect to novel sequences, as is the case in real-world applications, we created an additional test set for each ion channel, called the novel-test set, which has little or no similarities to the sequences in either the sequences of the train set or the test set. In the novel-test experiment, Multi-Branch-CNN performs significantly better than TML13, showing an improvement in accuracy of 6%, 14%, and 15% for sodium, potassium, and calcium channels, respectively. We confirmed the effectiveness of Multi-Branch-CNN by comparing it to the standard CNN method with one input branch (Single-Branch-CNN) and an ensemble method (TML13-Stack). To facilitate applications, the data sets, script files to reproduce the experiments, and the final predictive models are freely available at https://github.com/jieluyan/Multi-Branch-CNN.

## Introduction

Ion channels are a family of transmembrane proteins that form pores in the membrane. They regulate the influx and effiux of cations or anions in cells. Because of the important role that ion channels play in both excitable and nonexcitable tissues, they are attractive therapeutic targets for many diseases, including neurological disorders, cardiovascular and metabolic diseases, and cancers. Peptides that bind ion channels play an important role in regulating ion flux across the plasma membrane. They are molecules commonly used for pharmacological characterization of various ion channels and receptors [1, 2]. However, experimental methods to search for novel interacting peptides are expensive, so computational methods are needed [3].

While there are several computational works that address peptide prediction for ion channels, most of them use limited machine learning (ML) and feature encoding methods. Moreover, to our knowledge, there are no online prediction tools available and the public databases dedicated for ion channel interacting peptides are limited. The first computational work for ion channel peptide prediction was undertaken by Mei *et al* [4]. They compared 7 traditional ML methods, namely Random Forest (RF), naïve bayes multinomial (NBM), support vector machines (SVM), logistic regression (LR), Instance Based Learner (IBk), J48 and multilayer perceptron (MP) and investigated 2 different feature encoding methods, amino acid composition (AAC) and di-peptide composition (DPC) as well as their combinations after feature selection. Moreover, 3 RF models to identify inhibitory peptides of calcium, potassium, and sodium channels were developed and achieved a sensitivity of 60.00% for calcium channels, 71.90% for potassium channels, and 86.80% for sodium channels when evaluated by the jackknife test. However, neither the datasets nor the program developed in this work have been released. Later on, Lissbet et al. [5] established an interface of PPLK^+^C. As this work focused only on identifying inhibitory peptides for potassium channels, its applicability to other ion channels cannot be certain.

In the present work, we propose a deep learning method, Multi-Branch-CNN, for ion channel peptide prediction based on CNN. The idea behind the term “parallel” is that the predictor can accept and learn from multiple feature types. A particular challenge with ML of peptides is to determine, from the vast amount of different protein or peptide descriptors [6], which of them contain features that are highly correlated with the prediction task at hand. An exhaustive or heuristic search using feature selection methods is always computationally intensive and may not lead to an optimal model. Since each feature type contains a complete but different views of the peptides, all features in a feature type should be treated as one collection. Although not explicitly discussed in the literature, it is generally accepted that all features in a feature type would be included when it is selected for model building. However, standard feature selection methods consider each feature independently and select each feature individually during the selection process rather than considering the entire feature set. When multiple feature types are found to perform equally well in learning a particular dataset, as it is quite often the case [7], these features are usually combined to form a feature matrix as input to the learning engine [8]. Alternatively, each feature type may be used to train a separate model, and finally these different models are ensembled to produce a single prediction [9, 10]. With Multi-Branch-CNN, a set of feature types with good performance can be jointly included in the learning process. In a comparative study of Multi-Branch-CNN with thirteen traditional ML methods (TML13), Multi-Branch-CNN shows comparable performance in general test cases - test sequences that have moderate similarity to those in training sets. However, when novel sequences were tested - sequences that have very little or no similarities to sequences in the training sets - Multi-Branch-CNN shows a dramatic improvement in classification accuracy compared to TML13. Moreover, to ensure the effectiveness of Multi-Branch-CNN, its performance is compared to Single-Branch-CNN, which has only 1 CNN input branch and an ensemble method TML13-Stack. In this work, we also explore different ways of preparing negative data set and show that properly processed negative data is crucial for developing reliable prediction models.

## Materials and data preprocessing methods

### Dataset Preparation

#### Ion channel interacting peptides (positive data)

The known ion channel interacting peptides of sodium (referred to herein as Na-pep), potassium (K-pep), calcium (Ca-pep) channels were downloaded from the Swiss-Prot [11] database using the keywords: `“sodium channel impairing toxin” AND reviewed:yes; “potassium channel impairing toxin” AND revie wed:yes; “calcium channel impairing toxin” AND reviewed:yes.` For each dataset, we first removed the peptide sequences containing unnatural or arbitrary amino acids (B, J, O, U, Z, X). For each valid sequence, if the post-translational modification section was present, only the longest sequence segment specified in the *chain* or *peptide* subsection was extracted, i.e., the extent of a polypeptide chain or active peptide in the mature protein. Sequences longer than 100 amino acids (a.a.) were deleted and duplicated sequences were removed. Additional interacting peptides for potassium channels were downloaded from the Kalium database [12] and subjected to the same preprocessing steps as described above. Finally, we obtained 728 Na-pep, 580 K-pep, and 246 Ca-pep. The numbers of the collected sequences are listed in Table 1. All the preprocessed sequences of the three channels can be downloaded in the supplementary file, such that the corresponding negative sequences generated by CDHit negative data generation method. The sequence length distribution and AAC of these peptides are presented in Figure 1.

**Table 1.** Summary of the ion channel interacting peptides downloaded from public repositories.

| Peptide | From Swiss-Prot | From Kalium | Total Valid Sequences | Length |
| --- | --- | --- | --- | --- |
| Na-pep | 820 | - | 728 | 5 – 83 |
| K-pep | 646 | 344 | 580 | 5 – 83 |
| Ca-pep | 333 | - | 246 | 5 – 83 |

**Fig. 1.**
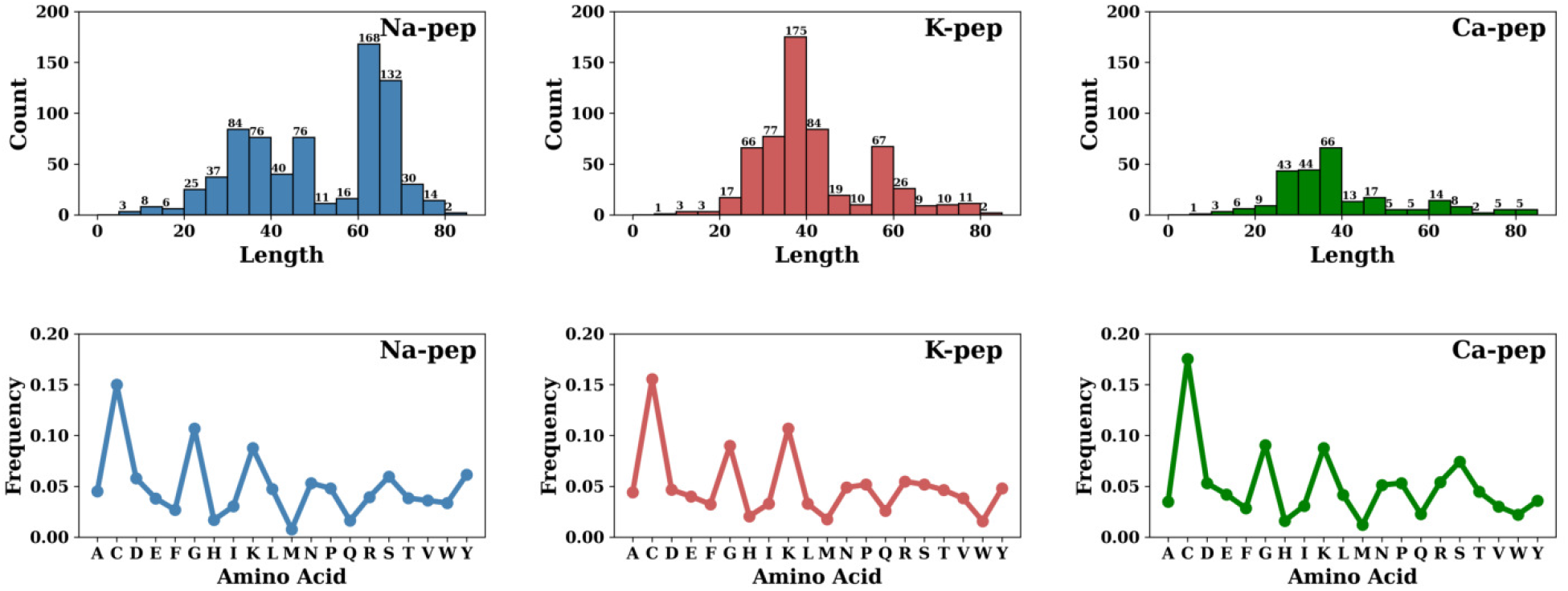
Sequence length distribution and AAC of sodium, potassium, and calcium ion channel interacting peptides.

#### Non-ion channel interacting peptides (negative data)

While positive data are usually reported in literature, negative data is scare. For the sake of supervised ML, we downloaded sequences without known channel interacting activities from Swiss-Prot [11] with the keywords `NOT goa:(“ion channel inhibitor activity [0008200]”) NOT goa:(“toxin activity [00 90729]”) NOT “ion channel impairing toxin” length:[1 TO 100]” reviewed:yes.` Following the same preprocessing procedure as for the positive data, we removed the peptide sequences containing unnatural amino acids and the duplicate peptides within the negative samples. At the same time, we also removed redundant sequences from the negative samples by comparing them with the positive datasets from all three channels. Finally, we obtained a total of 37,437 negative samples.

#### Dataset splitting: train, test, and novel-test

Conventionally, when developing a ML model, the entire dataset is divided into a train set and a test set by random splitting. In this way, the test set may contain a mixture of sequences with high and low similarity to the training sequences in unknown proportions, making it difficult to assess the generalizability of the developed models. To obtain a more reliable evaluation, we propose to use two separate test sets: First, a normal test set that contains sequences with general similarity to the train set (with at least 40% sequence similarity). Second, a novel-test set to estimate the predictive power of the model on truly novel sequences. These sequences have less than 40% sequence similarity with the train set and the test set.

The train, test and novel-test sets for all 3 ion channels (sodium, potassium, calcium) were created according to the workflow described in Figure 2. We first employed CD-HIT [13] to cluster sequences using a threshold of 0.4 and a word length of 2. For each cluster containing two or more member peptides, one peptide was randomly selected and used as test data. For peptides that could not be clustered, i.e., no similar sequences were found in the group, they were merged together to form the novel-test dataset. Finally, the unselected peptides were used as training data. We adjusted number of samples in the train and test sets to maintain a train:test ratio of 9:1. It should be noted that as CD-HIT only processes sequences whose length is greater than 10 a.a., to avoid data loss, these sequences were added back to the novel-test set.

**Fig. 2.**
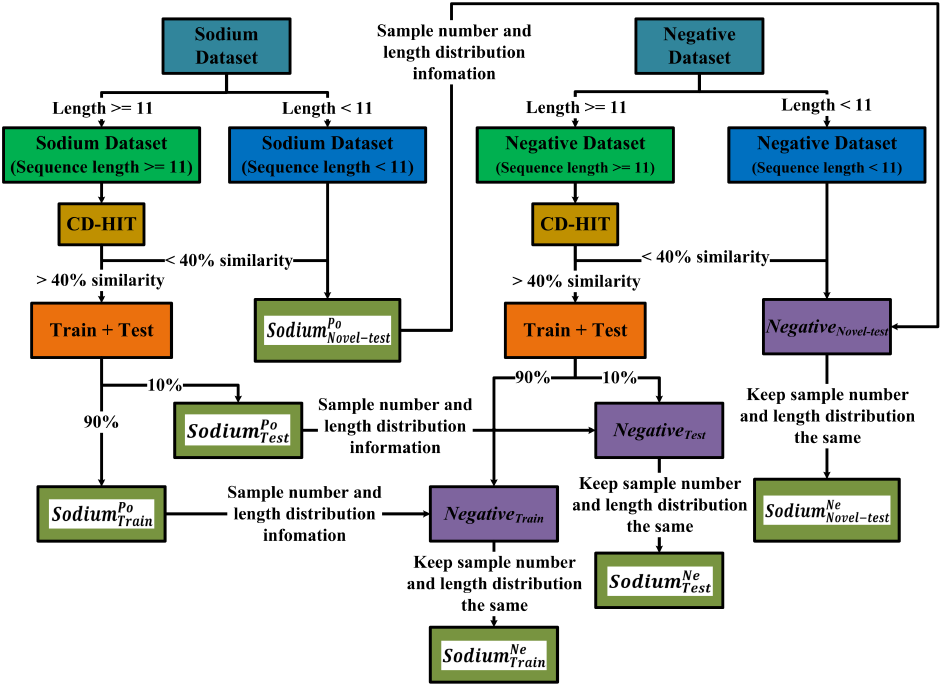
Workflow for creating the train, test, and novel-test sets using CDHit. The Na-Pep is shown here as an example. Positive dataset is indicated with the superscript *Po* and negative dataset with *Ne*. The role of each dataset in ML is indicated with the subscript *train, test*, or *noveltest*.

We tested four different approaches to generate negative datasets, which are described in detail below. The results of our comparative study showed that the best approach, which provided reliable models with stable performance was the use of CDHit. Here, we describe the generation of negative datasets using CDHit. The negative train, test and novel-test sets were created by drawing sequences from the clustered negative sequences in the same manner as for the positive sets. The number of sequences in the negative sets equals to the number of sequences in the positive sets, i.e., balance datasets were created. To avoid length bias, the peptides in a negative set were carefully selected so that the length distributions of the set matched the corresponding positive set. The final balanced train, test, and novel-test sets used for model construction and testing are listed in Table 2.

**Table 2.** Number of positive sequences in the train, test, and novel-test sets. The same number of negative sequences were created in each set using four different approaches: normal, CDHit, random, and shuffle.

| Peptide | Train | Test | Novel-test | Total |
| --- | --- | --- | --- | --- |
| Na-pep | 639 | 71 | 18 | 728 |
| K-pep | 499 | 55 | 26 | 580 |
| Ca-pep | 211 | 23 | 12 | 246 |

#### Four approaches to generate negative data

The number of negative samples far exceeds the number of positive samples. The imbalance between the positive and negative data is approximately 1:50, 1:65, and 1:150 for Na-pep, K-pep, and Ca-pep, respectively. Highly imbalanced data are notoriously difficult to train, leading to the creation of impractical predictive models that have high specificity but far too low sensitivity. To create balanced datasets that support the development of reliable predictive models, we explored four different approaches for generating the balanced negative train, test, and novel-test sets.

##### Normal

The negative datasets were generated by randomly selecting peptides from the negative peptide sequences downloaded from Swiss-Prot. In this process, the number and length distribution of the negative datasets matched those of the positive datasets deliberately. Redundant peptides in the negative sets to the positive sets were removed.

##### CDHit

The negative datasets were first divided into train, test and novel-test sets using CDHit, as described above. Then, for the train, test, and novel-test positive datasets of each channel (sodium, potassium, and calcium channels), we randomly selected the same number of peptides from the corresponding negative train, test, and novel-test sets and kept the length distribution between the corresponding negative set and positive set the same. Redundant peptides from the negative sets to the positive sets were removed. The procedure is shown in Figure 2.

##### Random

For each sequence in the positive datasets (train, test, and novel-test sets of sodium, potassium, and calcium ion channels), we generated a negative sequence of the same length as the template sequence by randomly mutating each residue in the template to any natural amino acids. In doing so, we assumed that the probability of a peptide with random sequence being an ion channel interacting peptide is very low.

##### Shuffle

Instead of mutating the residues in a sequence, we shuffied the amino acids in the sequence of a positive peptide to create a negative sequence. Again, it is assumed that the probability of a random sequence peptide being an ion channel interacting peptide is very low.

### Performance Metrics

We used accuracy (*ACC*), area under the curve of ROC (*AUC*), specificity (*Sp*), sensitivity (*Sn*), precision, *F1-*score, *Kappa*, matthews correlation coefficient (*MCC*) as metrics to measure the performance of predictive models.

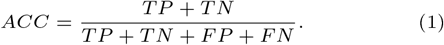

where *t* means the threshold to calculate *TP, TN, FP, FN*. The formula of *Sp* or true-negative rate (*TNR*) is presented as follows:

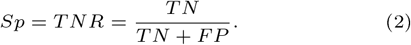

The formula of *Sn, recall, hit rate*, or true positive rate (*TPR*) is shown as follows:

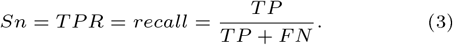

The formula of *AUC* is shown as follows:

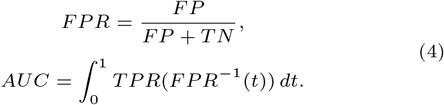

The formula of *precision*, or positive predictive value (*PPV*) is shown as follows:

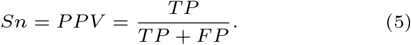

The formula of *F1-*score shown as follows:

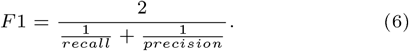

The formula of *Kappa* is shown as follows:

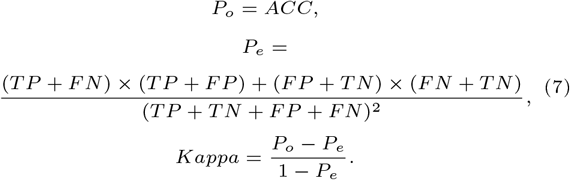

The formula of MCC is shown as follows:

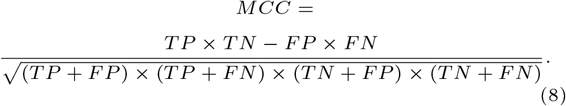

where *t* means the threshold to calculate *TP, TN, FP, FN*.

## Feature selection

The iFeature [6] package was used to numerically encode peptide sequences by generating 21 different feature types (such as AAC, CSKAAP, DPC, etc.) and 233 different PseKRAAC [14] feature groups. All feature types used in this study are listed in Table 3.

**Table 3.**
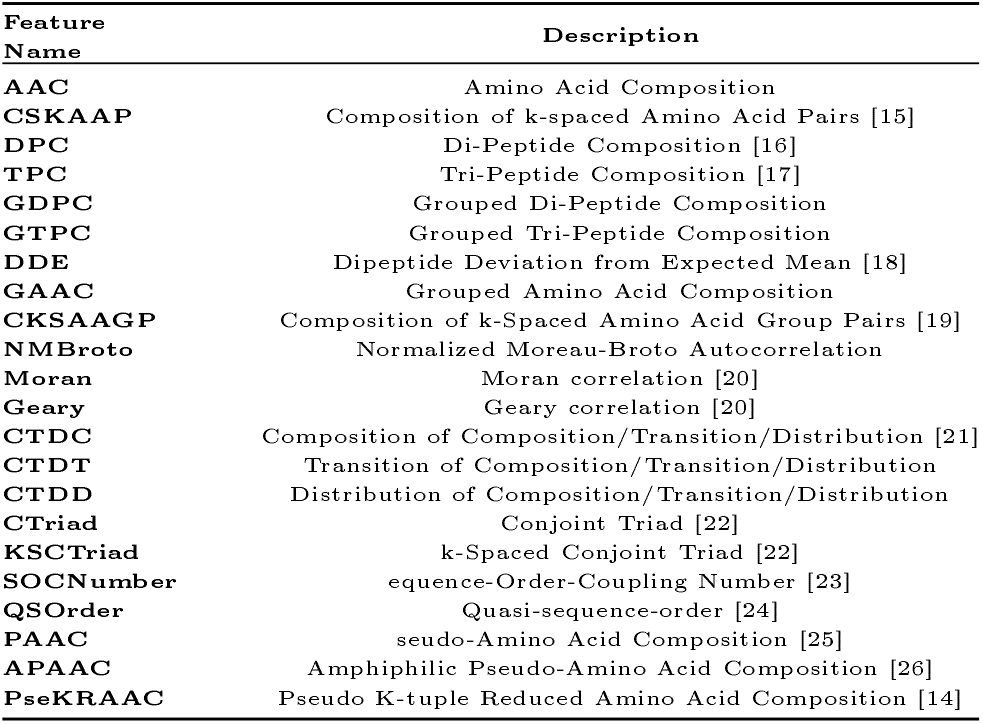
The feature types from the iFeature package used to encode peptide sequences

PseKRAAC [14] is a class of protein feature generation methods called Pseudo K-tuple Reduced Amino Acid Composition (PseKRAAC). PseKRAAC encodes peptide or protein features by implementing reduced amino acid alphabets (RAAC). By grouping related amino acids in different ways, this method attempts to reduce information redundancy in the sequence data and generate feature matrices with lower complexity and thus avoid overfitting. There are a total of 250 feature groups of PseKRAAC, however, RAAC with 20 groups were removed. RAAC that is too large (i.e., 20 groups) is the same as AAC, i.e., reduced encoding was not applied. After our preliminary test, only 233 feature groups of PseKRAAC were used further in the feature selection step. The following parameters were used in the PseKRAAC feature generation: *ktuple* = 2, gap-*λ* = 1, *gap* = 0, and *λ−*correlation = 4.

### Single feature type selection

Since there were many feature types under consideration (254 in total), it was computationally prohibitive to examine all combinations of feature types. However, to obtain the best optimal combination of feature types, we first examined the predictive power of each feature type and selected only those that performed reasonably good. Then, combinations of these feature types were tested to determine the optimal feature encoding approach.

The predictive performance of each feature type was measured using the TML13 metric (refer equation 9). In total, we performed 39,624 (254 feature types *×* 13 ML algorithms *×* 3 ion channel types *×* 4 negative dataset generation methods) experiments for single-feature-type selection.

### Best-*K* features combination selection

We computed the TML13 metrics of all best-*K* features combination and sorted them by average accuracy over all 3 channels with the CDHit data generation method. A best- *K* features combination is denoted as {best-1, best-2, …, best-*K*} (*K* ∈ {1, 2, …, 30}), where best-*i* represents the feature type of the *i*^*th*^ (*i* ∈ {1, 2, …, *K*}) highest average accuracy over all 3 channels with CDHit data generation method on the train sets. The top 30 single feature types are “type3Braac9”, “type7raac19”, “type8raac16”, “type7raac18”, “type1raac18”, “type10raac18”, “type8raac14”, “CKSAAP”, “DDE”, “type7raac17”, “type11raac10”, “type12raac16”, “type12raac17”, “type8raac13”, “KSCTriad”, “type7raac16”, “type1raac19”, “type10raac19”, “type11raac18”, “type9raac16”, “type14raac18”, “type9raac15”, “type14raac17”, “type11raac16”, “AAC”, “type3Braac12”, “type7raac14”, “type12raac9”, “type14raac15”, and “type12raac11”, where ‘type*X*raac*Y* ‘ indicates the type *X* of PseKRAAC with the RAAC and *Y* number of clusters.

## Designed methods

### TML13 method

To make the predictive performance in the feature selection and to compare with Multi-Branch-CNN method, TML13 method was used, consisting of 13 traditional ML algorithms (listed in Table 4). The predictive performance was measured using the TML13 metric:

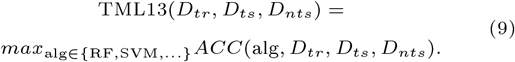

where *D*_*tr*_, *D*_*ts*_, *D*_*nts*_ are the train, test, and novel-test feature matrix datasets for sodium, potassium, or calcium ion channel that was generated using one of the four negative data generation methods. This metric is a triplet of three values [*ACC*_*tr*_, *ACC*_*ts*_, *ACC*_*nts*_] where *ACC*_*tr*_ is the highest accuracy in 5-fold cross-validation (CV) of *D*_*tr*_; *ACC*_*ts*_ and *ACC*_*nts*_ are the test and novel-test accuracies on *D*_*ts*_ and *D*_*nts*_, respectively, using the optimal model according to CV.

**Table 4.** 13 traditional ML algorithms of TML13.

| Algorithm | Description |
| --- | --- |
| **RF** | Random Forest [27] |
| **SVM** | Support Vector Machine [28] |
| **Ada** | Ada boost classifier [29] |
| **LightGBM** | Light Gradient Boosting Machine [30] |
| **GBC** | Gradient Boosting Classifier [31] |
| **ET** | Extra Trees classifier [32] |
| **LDA** | linear discriminant analysis [33] |
| **Ridge** | Ridge classifier [34] |
| **QDA** | Quadratic Discriminant Analysis [35] |
| **DT** | Decision Tree classifier [36] |
| **LR** | Logistic Regression [37] |
| **NB** | Naïve Bayes [38] |
| **KNN** | K Nearest Neighbour [39] |

### TML13-Stack method

In recent papers, ensemble method has been proposed as an effective method in peptide prediction methods [9, 10]. Therefore, a form of ensemble algorithm called TML13-Stack was implemented as a comparison. TML13-Stack is a two level method, it first sorts the 13 algorithms (shown in Table 4) by the average of 5-fold cross-validation accuracy on the training sets.

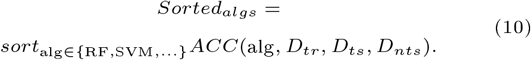

where the 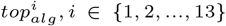 represents the *i*^*th*^ highest *ACC*_*tr*_ algorithm name, 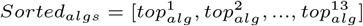 is a list of sorted algorithms. From this, with the predictions of the 2^*nd*^ highest to the 13^*th*^ highest algorithms 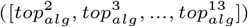 as the input to the second layer (stack layer), 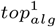 can be applied as the classification algorithm to make the final prediction. The first layer is:

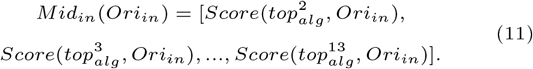

where *Ori*_*in*_ signifies the input set, *Mid*_*in*_ is the input of the stack layer, and the 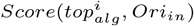 represents the predicted score of the model developed by 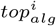 with *Ori*_*in*_ as the inputs. The stack layer is:

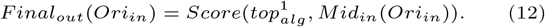

where *Final*_*out*_ corresponds to the predicted score of the stack layer. The predictive performance of each feature type was measured using the TML13-Stack metric:

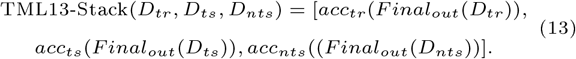

where *acc*_*tr*_ represents the average 5-fold corss-validation accuracy calculation function of the train set, *acc*_*ts*_ and *acc*_*nts*_ are defined as the functions to calculate the accuracies of the test or the novel-test sets, respectively.

### Proposed Multi-Branch-CNN Method

When predicting peptides that interact with ion channels, traditional ML methods performed quite well. However, an interesting phenomenon we observed is that different feature types could give very similar prediction performances, which makes it difficult to accurately select the feature subsets. The idea of the method proposed here is to exploit the predictive ability of different feature types without the need for exhaustive feature selection. By training individual CNN models based on the selected feature types, we combined the independent predictions of all CNN models to obtain a better combined prediction. The training procedure was designed to train the models in parallel, hence giving the name “Multi-Branch-CNN”.

### CNN architecture

As shown in Figure 3, the Multi-Branch-CNN model consisted of *K*-CNN models arranged in parallel, where *K* is the number of feature types selected for the learning task at hand. The input `Best`-*i*, where *i ∈* {1, 2, ..., *K*}, represents best-*i* of the best-*K* features combination selected in the feature selection stage. To treat the features as images and input them in parallel into our CNN architecture, we convert each feature group into a square matrix with dimension 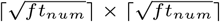, where *ft*_*num*_ is defined as the feature number of the corresponding feature group. Note that we add zeros to the end of the feature groups if the size of the square matrix is larger than the feature number. For the prediction of ion channel interacting peptides, a combination of *K* (18 in this paper, refer to Figure 6) feature types proved to be better than other combinations. Here, each CNN component model was comprised of two convolutional layers followed by a flattened layer and a 64-node dense layer. All convolutional layers consisted of a stack normalization part, a 2D convolutional part with filters whose size is 3 *×* 3 and equal padding, and a max-pooling part with 2 *×* 2 and equal padding. All activation functions were set to relu except for the output layer, which was `sigmoid`.

**Fig. 3.**
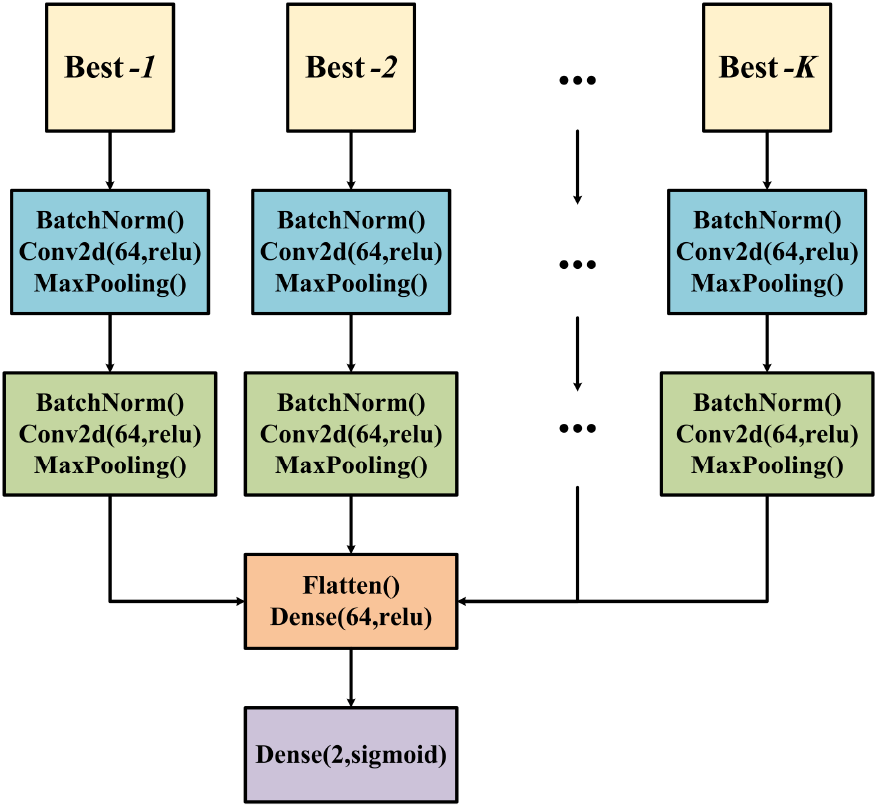
The proposed Multi-Branch-CNN model with optimal parameters.

### CNN parameter tuning

To find the best CNN architecture, we tested the parameters of 1 and 2 convolutional layers by setting the dropout rate as 0, 0.1, 0.2, 0.3, 0.4 or 0.5, the filter number of each layer as 16, 32, 64, 96 or 128, and the epoch as 1, 2, 3, …, or 500. The parameter search was performed by grid search and the results were sorted by validation accuracy. In total, there were 2 *×* 6 *×* 5 *×* 500 experiments to find the best hyperparameters of the final Multi-Branch-CNN model. The final hyperparameters are as follows: CNN layer = 2, filter = 64, dropout = 0.4, and epoch = 187. To obtain a stable performance in the prediction model, we manually tested the use of an early stopping criterion over different settings of the parameters patience and monitor, as well as training using different learning rates, decay rates and numbers of decay steps. Finally, we found that early stopping monitored with “loss”, via a learning rate of 0.0001 and patience of 20 were the best combination. We also determined that the proposed CNN architecture with early stopping was better when the loss was set to 0. For both Na-pep and K-pep, predictions, when using a decay rate of 0.9 and decay steps of 100, the proposed architecture performed the best. As for Ca-pep predictions, a decay rate of 0.86 and decay steps of 50 for the proposed architecture performed the best.

### Single-Branch-CNN method

In order to prove the effectiveness of Multi-Branch-CNN, a 1-CNN model was implemented for comparison called Single-Branch-CNN. Single-Branch-CNN shares the same parameters with Multi-Branch-CNN, but only consists of an 1-CNN architecture with the best-*K* features combination selected in the feature selection stage. The input of Single-Branch-CNN is a square matrix with dimension 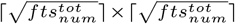, where 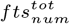 represents the total feature number of all the best-*K* features combination.

## Results

### Single feature selection

#### Comparative study of negative data generation methods

To get an overview of the predictive power of all single feature type models using traditional ML algorithms, we plotted the distribution of the training, testing, and novel-testing results of all single feature models on 3 channels of 4 data generation methods in Figure 4. As shown in Figure 4 (A), the Na-pep models performed better than the K-pep and Ca-pep models in both the training and test sets. However, in the novel-test set, the Ca-pep models performed the best. We noted that the training set of K-pep is the largest among the three, while Ca-pep’s is the smallest. It seems to suggest that the size of the training set not only plays a role in the prediction of similar peptides (test sequences), but also affects, perhaps negatively, the accuracy of the prediction of dissimilar peptides (novel test sequences).

**Fig. 4.**
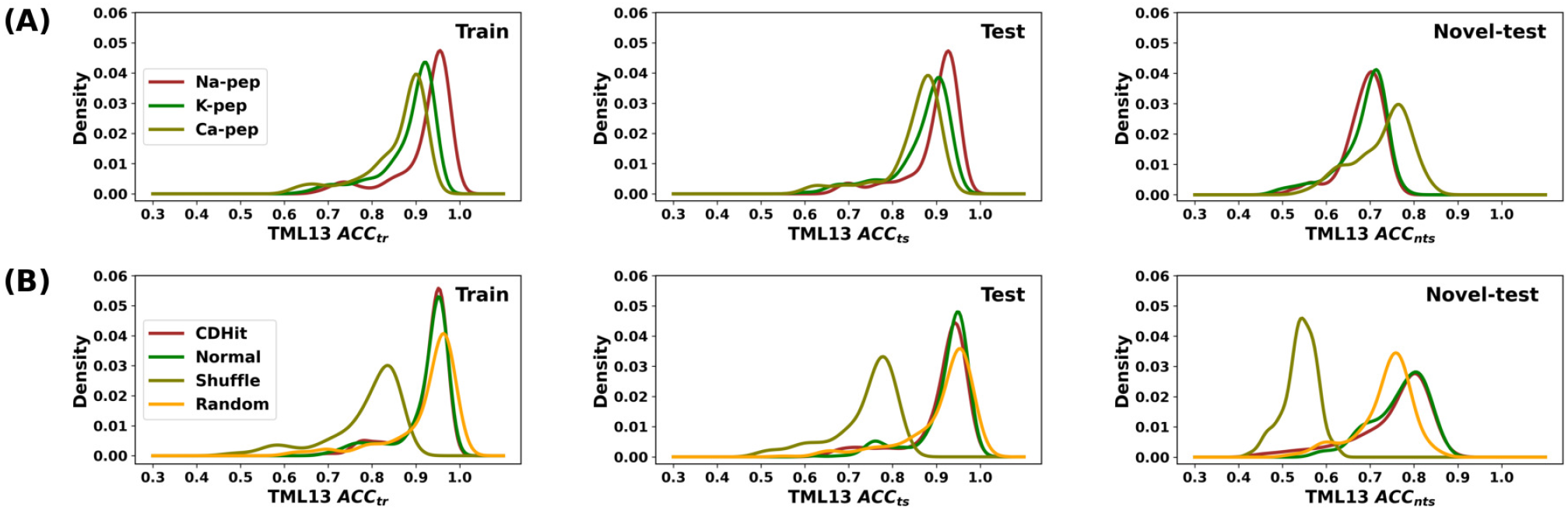
Distribution of the mean TML13 scores (*ACC*_*tr*_, *ACC*_*ts*_, *ACC*_*nts*_) of (A) Na-Pep, K-Pep, Ca-Pep averaged over 4 negative data generation methods, (B) four negative data generation methods averaged over 3 channel peptides. Each distribution was constructed from 254 TML13 scores, each for one single feature type model. For visual clarity, the distribution was smoothed by Gaussian kernel [40].

Figure 4 (B) compares the performance of models constructed using different negative dataset generation methods. It can be observed that Normal and CDHit perform better than Shuffie and Random in most single feature encoding models. Moreover, a comparison of the density curve of CDHit with that of Normal shows that the former is smoother while the latter is more fluctuant. A possible explanation for this is that the data samples in CDHit are more evenly distributed, as very similar sequences have been filtered out to ensure that they are not overrepresented in the dataset. Therefore, we conclude that using CDHit to create a non-redundant dataset for ML is a beneficial procedure. Henceforth, CDHit was used to process the final negative dataset, which was subsequently used to create the production models.

#### Comparative study of traditional ML methods

It would also be interesting to know which traditional ML algorithm performed best. Figure 5 shows the heatmap of the average performance of each algorithm for predicting the peptides of each channel using different training sets. Each value was averaged over 254 single feature types (21 feature types and 233 PseKRAAC feature groups). It can be observed that ET, LightGBM, RF, GBC, LR are the top five performing algorithms, regardless of the channel types or negative data generation method used. In all cases, QDA performed the worst.

**Fig. 5.**
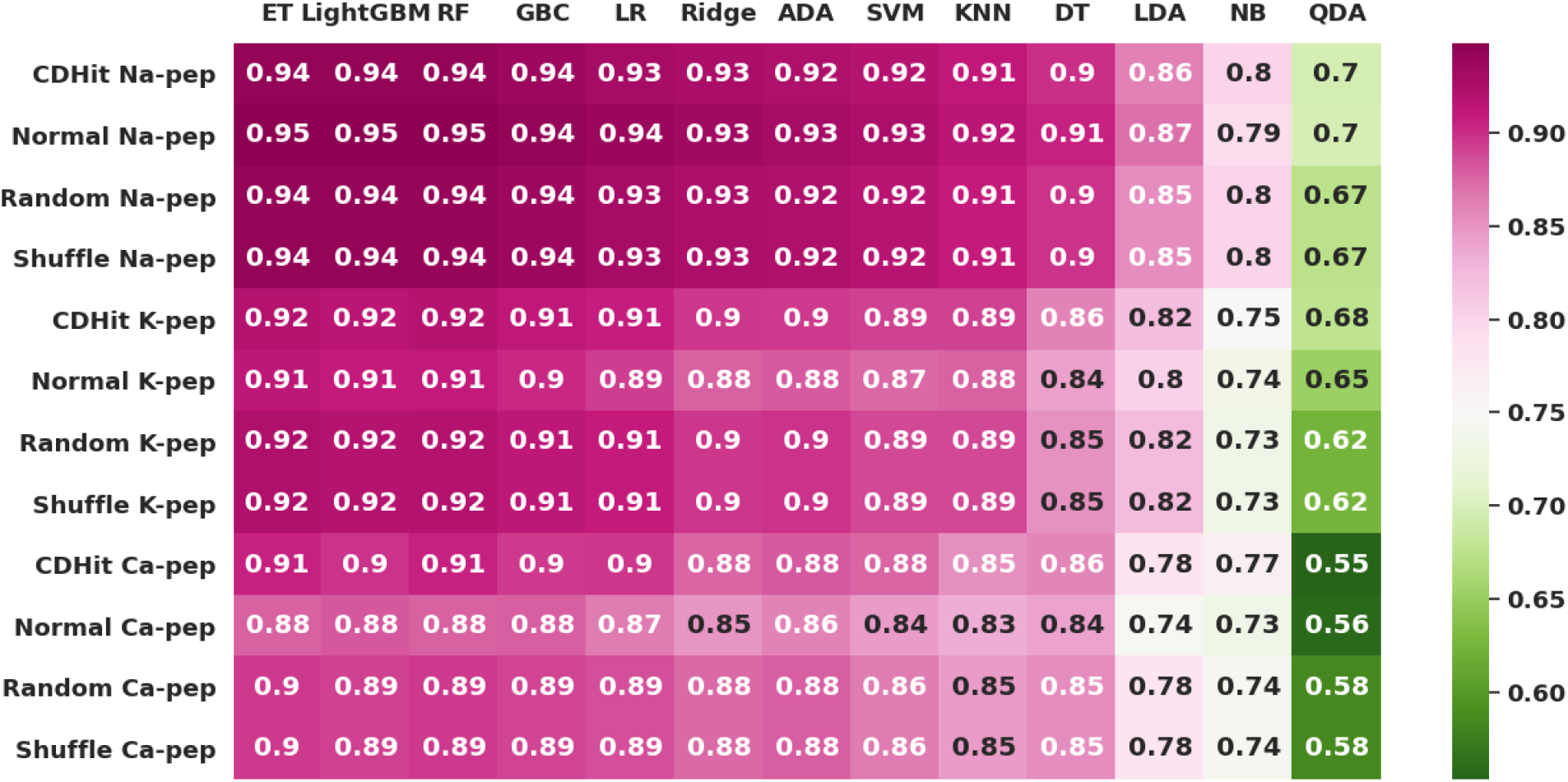
The average 5-fold cross-validation accuracy of each TML algorithm on Na-pep, K-pep, and Ca-pep predictions using the four di erent training sets.

### Best-*K* features combination selection

To determine how many feature types were needed to create the optimal models for channel peptide prediction, we evaluated the combinations of best 1, best 2, best 3, etc., up to best 30 feature types, where the rank of each feature type was the TML13 *ACC*_*tr*_ score using the CDHit datasets. As shown in Figure 6, the best-18 features combination achieved the highest average cross-validation accuracy in the training sets of all three channels. These best-18 features are “type3Braac9”, “type7raac19”, “type8raac16”, “type7raac18”, “type1raac18”, “type10raac18”, “type8raac14”, “CKSAAP”, “DDE”, “type7raac17”, “type11raac10”, “type12raac16”, “type12raac17”, “type8raac13”, “KSCTriad”, “type7raac16”, “type1raac19”, and “type10raac19”. It is interesting that most of the best feature types are from the PseKRAAC feature groups, differing by their types and number of reduced amino acid clusters. The best-18 features are listed in Table 5. From Figure 6, some test accuracies can reach the average cross-validation accuracy of the training set, but novel-test accuracy performs much lower than that of the test accuracy and the average cross-validation accuracy of the training set, which shows the same trends in the single feature selection plot in Figure 4.

**Fig. 6.**
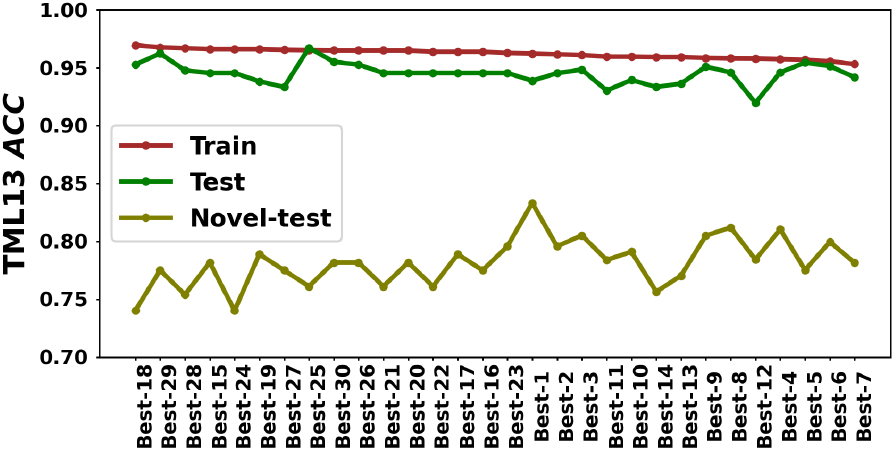
Cross-validation performance of best-1 to best-30 feature types combination in TML13 using the CDHit datasets.

**Table 5.** The best-18 feature types and their number of features.

| Feautre Groups | Type | RAAC Type | Feature number |
| --- | --- | --- | --- |
| PseKRAAC | 3B | 9 | 81 |
| PseKRAAC | 7 | 19 | 361 |
| PseKRAAC | 8 | 16 | 256 |
| PseKRAAC | 7 | 18 | 324 |
| PseKRAAC | 1 | 18 | 324 |
| PseKRAAC | 10 | 18 | 324 |
| PseKRAAC | 8 | 14 | 196 |
| CKSAAP | - | - | 800 |
| DDE | - | - | 400 |
| PseKRAAC | 7 | 17 | 289 |
| PseKRAAC | 11 | 10 | 100 |
| PseKRAAC | 12 | 16 | 256 |
| PseKRAAC | 12 | 17 | 289 |
| PseKRAAC | 8 | 13 | 169 |
| KSCTriad | - | - | 686 |
| PseKRAAC | 7 | 16 | 256 |
| PseKRAAC | 1 | 19 | 361 |
| PseKRAAC | 10 | 19 | 361 |

#### Comparison between TML13 and Multi-Branch-CNN methods

Using the best-18 features combination and the CDHit training sets, we created the final optimal models for Na-pep, K-pep, and Ca-pep. The models were tested against the test sets and novel test sets of all three channel peptides. We should point out that a test set contains sequences that were selected from each sequence cluster in the training sets, while a novel-test set contains sequences with little or no sequence similarity to sequences in the training sets and are truly unseen sequences. The training and testing experiment was repeated 5 times and the test accuracies were averaged.

The performance comparison between TML13 and the Multi-Branch-CNN methods on Na-pep, K-pep, and Ca-pep are shown in Tables 6, 7 and 8, respectively. However, on novel-test predictions, Multi-Branch-CNN significantly outperforms TML13 by improving the prediction accuracy by 6%, 14%, and 15% for Na-pep, K-pep, and Ca-pep, respectively. As for stability, TML13 models showed a more stable result with a standard deviation close to zero after repeating the experiment. 0n the other hand, the performance variation of the Multi-Branch-CNN models is also low, ranging from 0.01 to 0.035. 0verall, Multi-Branch-CNN predicts a more stable result than TML13 for a majority of the metrics under the general test sequences, while it predicts with a 6 - 15% higher accuracy for truly novel sequences. This result is remarkable, suggesting that Parellel-CNN is more capable to discover novel ion channel interacting peptides than other predictors.

**Table 6.** Comparative performance of TML13, and Multi-Branch-CNN on Na-pep predictions.

| Test | ACC | Sp | Sn | Prec | F1 | Kappa | MCC |
| --- | --- | --- | --- | --- | --- | --- | --- |
| TML13 | 0.951 | 0.944 | 0.958 | 0.944 | 0.951 | 0.901 | 0.901 |
| Multi-Branch-CNN | 0.98 | 0.966 | 0.994 | 0.967 | 0.981 | 0.961 | 0.961 |
| Novel-test | ACC | Sp | Sn | Prec | F1 | Kappa | MCC |
| TML13 | 0.714 | 0.941 | 0.5 | 0.9 | 0.643 | 0.436 | 0.488 |
| Multi-Branch-CNN | 0.760 | 0.941 | 0.589 | 0.9139 | 0.7159 | 0.525 | 0.563 |

**Table 7.** Comparative performance of TML13 and Multi-Branch-CNN on K-pep predictions.

| Test | ACC | Sp | Sn | Prec | F1 | Kappa | MCC |
| --- | --- | --- | --- | --- | --- | --- | --- |
| TML13 | 0.973 | 0.945 | 1 | 0.948 | 0.974 | 0.946 | 0.947 |
| Multi-Branch-CNN | 0.953 | 0.9209 | 0.985 | 0.9259 | 0.9549 | 0.9059 | 0.908 |
| Novel-test | ACC | Sp | Sn | Prec | F1 | Kappa | MCC |
| TML13 | 0.714 | 0.870 | 0.577 | 0.833 | 0.682 | 0.438 | 0.462 |
| Multi-Branch-CNN | 0.816 | 0.835 | 0.8 | 0.846 | 0.822 | 0.632 | 0.634 |

**Table 8.** Comparative performance of TML13 and Multi-Branch-CNN on Ca-pep predictions.

| Test | ACC | Sp | Sn | Prec | F1 | Kappa | MCC |
| --- | --- | --- | --- | --- | --- | --- | --- |
| TML13 | 0.934 | 0.913 | 0.957 | 0.917 | 0.936 | 0.87 | 0.87 |
| Multi-Branch-CNN | 0.939 | 0.896 | 0.983 | 0.905 | 0.942 | 0.878 | 0.882 |
| Novel-test | ACC | Sp | Sn | Prec | F1 | Kappa | MCC |
| TML13 | 0.792 | 0.75 | 0.833 | 0.769 | 0.8 | 0.583 | 0.585 |
| Multi-Branch-CNN | 0.908 | 0.817 | 1 | 0.847 | 0.917 | 0.817 | 0.832 |

We attempted to compare Multi-Branch-CNN with previous methods published in the literature. Unfortunately, the only two computational studies of ion channel peptides were Mei et *al*. [4] and Lissbet *et al*. [5], but neither their program codes nor datasets were available, making it difficult to make a fair comparison with them. According to the performance results published in Lissbet *et al*. [5], the potassium channel peptide prediction by PPLK^+^ C method has an accuracy of 0.94 and the method of Mei *et al*. [4] has 0.86, while our Multi-Branch-CNN method, with an updated dataset, achieves a higher accuracy than these two.

#### Further comparative study with TML13-Stack and Single-Branch-CNN methods

Although the Multi-Branch-CNN method performs better than the TML13 method from Table 6, 7, and 8, it is not strong enough to prove that it is better than an ensemble method or a normal CNN method with only 1 input branch. Therefore, Single-Branch-CNN, a Multi-Branch-CNN architecture that inputs the best-18 features combination (shown in Figure 5) as only 1 square matrix, was developed to assess whether CNN branches with multiple inputs performs better than CNN branches with only one input. Also, TML13-Stack was developed to test whether the Multi-Branch-CNN performs better than an ensemble method. It should be noted that the best-18 features combination selected in the feature selection stage was utilized as the feature encoding method for all the four methods (TML13, TML13-Stack, Single-Branch-CNN, and Multi-Branch-CNN).

Figure 7 shows a visualization of the average accuracy of 5 times repetitions by the 4 methods, namely TML13, TML13-Stack, Single-Branch-CNN, and Multi-Branch-CNN for 3 channels and the 4 methods’ average of the 3 channels on the test and novel-test sets. For the test sets, Multi-Branch-CNN performs on par with all the other 3 methods (TML13, TML13-Stack, and Single-Branch-CNN) for all 3 channels and slightly better than the other 3 methods when it comes to the average. The TML13-Stack method performs the worst on the test sets for all 3 channels. The Single-Branch-CNN method performs worse than the TML13 method for K-pep, but better than the TML13 method for Na-pep and Ca-pep on the test sets. As for the novel-test sets, Multi-Branch-CNN performs best and better than the second ranked methods for K-pep and Ca-pep, as well as the average of the 3 channels (see the average bar). Single-Branch-CNN performs best and slightly better than Multi-Branch-CNN for Na-pep on the novel-test sets. TML13-Stack performed terribly for Na-pep and K-pep on the novel-test sets. Moreover, TML13 and TML13-Stack attained a significantly worse result than the 2 CNN methods on the novel-test sets.

**Fig. 7.**
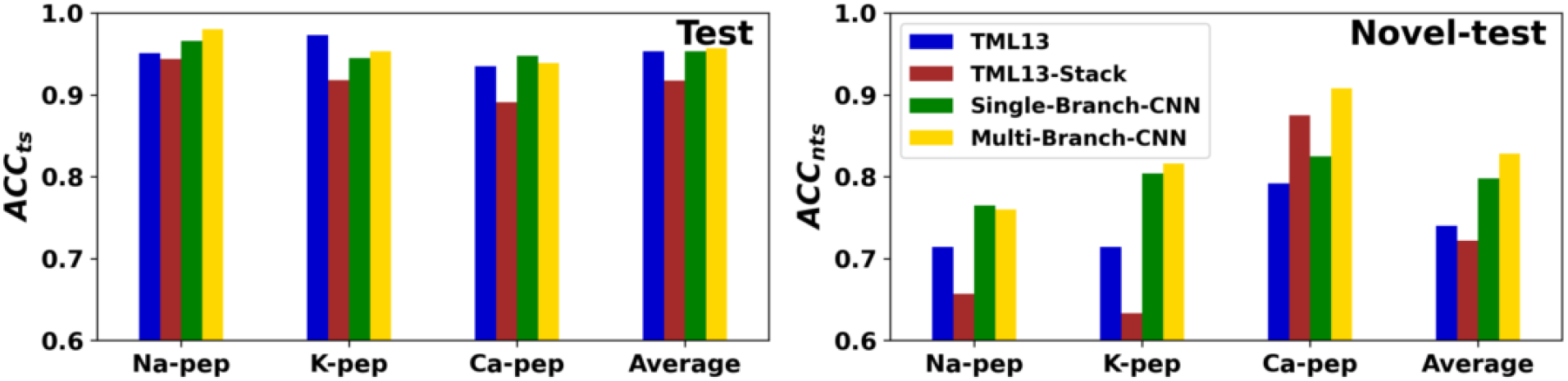
Comparative accuracy of TML13, TML13-Stack, Single-Branch-CNN and Multi-Branch-CNN on test left and novel-test right sets of Na-pep, K-pep, and Ca-pep predictions.

## Conclusion

In this work, we proposed a parallel CNN method (Multi-Branch-CNN) to predict sodium, potassium, and calcium ion channel interacting peptides. We extensively tested a large number of feature encoding methods (254 in total) and prioritized them for model construction. Four different methods for generating negative data was discussed, where we discovered that using the method CDHit is a reliable preprocessing step to generate a set of negative sequences that facilitates predictive modeling. To test the predictive performance of our method on truly unseen sequences, which is often the case in real-world applications, we created novel-test sets with the sequences having little or no similarity to the sequences in the training and/or test sets. Based on our extensive experiments, we confirmed that Multi-Branch-CNN is able to maintain a high prediction performance not only on general test sets, but can also achieve a comparatively good performance compared to traditional ML methods (TML13) by showing a 6-15% improvement in accuracy for predicting all three channel peptides. We also established that the Multi-Branch-CNN method performs better than an ensemble algorithm (TML13-Stacked) and a normal CNN method with only 1 input branch (Single-Branch-CNN).

## Data and Software Availability

All peptide data used in this study were downloaded from Swiss-Prot [11] and the Kalium database [12]. The data can be download from supplementary, and script files for reproducing the experiments can be downloaded from https://github.com/jieluyan/Multi-Branch-CNN. The final predictive models are also available at the above link. The implementation was done using Python 3.6.13, pycaret 2.3.1 [41], and tensorflow 2.2.0 [42].

## Author’s contributions

S.W.I.S. and H.F.K. conceived the study. J.L.Y. designed the methods, conducted the experiments, analyzed the results, and drafted the manuscript. S.W.I.S., M.Z. and B.Z. supervised the project. S.W.I.S., H.F.K., M.Z. and B.Z. finalized the manuscript. All the authors read and approved the final manuscript. S.W.I.S., H.F.K., B.Z. acquired project funding.

## Acknowledgements

This project was supported by University of Macau (grant no. MYRG2019-00098-FST) and the Science and Technology Department Fund of Macau SAR (File no. 0010/2021/AFJ).

**Key Points**

- We propose Multi-Branch-CNN, a parallel convolutional neural networks (CNNs) method for identifying three types of ion channel peptide binders (sodium, potassium, and calcium).
- Our Multi-Branch-CNN performs best on both of the test sets and the novel-test sets compared with TML13, TML13-Stack, and Single-Branch-CNN over the average accuracy of the three channels.
- We propose a new test set called novel-test set, which has little or no similarities to the sequences in either the sequences of the train set or the test set, to simulate the novel sequences in the real-world applications.
- We tested four different approaches to generate negative datasets, and finally, CDHit was selected by providing reliable models with stable performance.

